# Gemcitabine therapeutically disrupts essential SIRT1-mediated p53 repression in Atypical Teratoid/Rhabdoid Tumors

**DOI:** 10.1101/2023.09.02.556021

**Authors:** Dennis S. Metselaar, Michaël H. Meel, Joshua R. Goulding, Piotr Waranecki, Mark C. de Gooijer, Marjolein Breur, Jan Koster, Sophie E.M. Veldhuijzen van Zanten, Marianna Bugiani, Pieter Wesseling, Gertjan J.L. Kaspers, Esther Hulleman

**Author notes:** Corresponding author: Dr. Esther Hulleman Heidelberglaan 25 3584 CS, Utrecht. **Funding:** All research described has been funded by the Children Cancer-free Foundation (KIKA) (https://www.kika.nl/), a non-profit organization that funds pediatric oncology research. Authorship: D.S.M. and E.H. conceived and designed the project. D.S.M., M.H.M., J.R.G., M.C.G., and P.Wa. developed and validated the *in vitro* and *in vivo* models used in the study. D.S.M., M.H.M., J.R.G., and P.Wa. performed the functional *in vitro* and western blotting experiments. D.S.M., M.C.G., M.Br., and M.Bu. performed the immunohistochemistry experiments. D.S.M., M.H.M., and P.Wa designed and performed the *in vivo* trials. S.E.M.V.Z. performed and analyzed clinical imaging. P.We. acquired patient material and performed and validated methylation profiling. J.K. provided bioinformatical and statistical expertise and support. M.Bu and P.We provided pathological expertise and support. G.J.L.K. and E.H. acquired funding and supervised the study. All authors contributed to writing the manuscript.

## Abstract

**Background:** Atypical Teratoid/Rhabdoid Tumors (ATRT) are highly malignant embryonal tumors of the central nervous system with a dismal prognosis. Despite recent advances in understanding the molecular characteristics and subclasses of these tumors, effective therapeutic options remain scarce.

**Methods:** In this study, we developed and validated a novel patient-derived ATRT culture and xenograft model, which we used alongside a panel of other primary ATRT models for large-scale drug discovery assays. The identified hits were mechanistically and therapeutically investigated using an array of molecular assays and two orthotopic xenograft murine models.

**Results:** We found that ATRT are selectively sensitive to the nucleoside analogue gemcitabine, with additional efficacy in Sonic Hedgehog (SHH)-subtype ATRT. Gene expression profiles and protein analyses indicated that gemcitabine treatment causes degradation of Sirtuin 1 (SIRT1), resulting in cell death through activation of NF-kB and p53. Furthermore, we discovered that gemcitabine-induced loss of SIRT1 results in a nucleus-to-cytoplasm translocation of the SHH signaling activator GLI2, explaining the additional gemcitabine sensitivity in SHH-subtype ATRT. Treatment of SHH-subgroup ATRT xenograft-bearing mice with gemcitabine resulted in a >30% increase in median survival (p<0.005, log-rank test) and yielded long-term survivors in two independent patient-derived xenograft models.

**Conclusions:** These findings demonstrate that ATRT are highly sensitive to gemcitabine treatment, and we propose that gemcitabine may form part of a future multimodal treatment strategy for ATRT.

**Key points:**

- ATRT are specifically sensitive to gemcitabine treatment
- SIRT1 may serve as a novel therapeutic target in ATRT
- Gemcitabine should be considered for clinical use in ATRT patients

**Importance of the study:** Atypical Teratoid/Rhabdoid Tumors (ATRT) are highly malignant pediatric brain tumors with a 5-year survival of merely 30%, for which effective treatment options are limited. In this study, we propose a potential novel treatment strategy for ATRT patients. We show that ATRT are highly sensitive to the chemotherapeutic gemcitabine, that takes advantage of ATRT-specific SIRT1 overexpression and disrupts p53 suppression and hedgehog signaling. Importantly, we show that gemcitabine significantly prolongs survival of ATRT patient-derived xenograft models, prolonging survival by over 30%. This effect was achieved using gemcitabine concentrations that are achievable in human brain and well-tolerated in pediatric patients. As such, gemcitabine could be readily incorporated into clinical treatment protocols and expand the still very limited therapeutic options for ATRT-patients.

## Introduction

Atypical teratoid rhabdoid tumors (ATRT) are highly malignant embryonal tumors of the central nervous system (CNS), often found in infants and young children. ATRT have historically been considered incurable tumors, and even though outcomes have recently improved slightly due to multimodal therapy, standardized treatment regimens are often absent and over 70% of all ATRT patients succumb within 5-years post-diagnosis.^1,2^ ATRT are characterized by bi-allelic loss of function of the tumor-suppressor SMARCB1, or in rare cases SMARCA4, which are core subunits of the SWI/SNF chromatin remodeling complex.^3^ Genome-wide methylation and RNA-sequencing experiments have distinguished three distinct molecular subgroups of ATRT, all typically characterized by low mutational burden and bland chromosomal copy number variation profiles: Sonic Hedgehog (SHH), Myc (MYC), and Tyrosinase (TYR).^4,5^ This subdivision of ATRT into specific molecular subgroups is an important step in understanding these clinically heterogenic malignancies, enabling researchers and clinicians to seek novel tailored therapeutic strategies.

SHH-ATRT is the most common of the three subgroups and is characterized by activation of the SHH signaling pathway, as demonstrated by strong overexpression and activation of GLI2 and MYCN.^4,6^ SHH-ATRT harbor a compact, hypermethylated chromatin structure with a strong epigenetically dysregulated expression profile, including overexpression of EHMT2 (G9a), EZH2, and several bromodomain containing proteins (BRDs). ^4,5^ Aside from the differences between each subgroup, most ATRT share the similarity of having functional, but suppressed, tumor-suppressor genes, like p53.^7,8^

In this study, we developed a new SHH-subtype ATRT culture and xenograft model and found, in multiple primary models, that ATRT are specifically sensitive to treatment with the clinically registered chemotherapeutic agent gemcitabine. Using RNA-sequencing, protein analysis, knockdown, and knockout experiments, we established that this ATRT-specific gemcitabine-sensitivity results from suppression of SIRT1, which causes p53 activation. We validated our findings in ATRT xenograft mouse models that showed significant increases in survival upon gemcitabine treatment. Furthermore, we discovered additional toxicity of gemcitabine treatment in SHH-subgroup ATRT, caused by gemcitabine induced inhibition of the SHH-signaling pathway. The results of our study warrant clinical investigation of gemcitabine as a potential therapy for ATRT patients.

## Materials and methods

### Patient material

Tumor tissue was obtained through surgical resection from an 11-year old patient that underwent surgery in the Amsterdam University Medical Center (Amsterdam, the Netherlands) for a tumor of which the clinical pathological diagnosis was ATRT. All patient material was collected according to national and institutional guidelines and in accordance with the declaration of Helsinki and put into culture as described previously.^9^

### Cell cultures

CHLA-ATRT-02 (SHH-ATRT), CHLA-ATRT-04 (SHH-ATRT), CHLA-ATRT-05 (SHH-ATRT) and CHLA-ATRT-06 (MYC-ATRT) cultures were obtained from the American Type Culture Collection (ATCC). VUMC-ATRT-01 (SHH-ATRT)^10^ and VUMC-ATRT-03 (SHH-ATRT) were established from tumor tissue obtained by surgical resection at the Amsterdam UMC (Amsterdam, the Netherlands) as described previously.^9^ CHLA-ATRT-266 (MYC-ATRT) was a kind gift from Dr Alonso (University of Navarra, Pamplona, Spain).

Non-ATRT pediatric glioma models VUMC-DIPG-A, VUMC-DIPG-F, VUMC-DIPG-10, VUMC-DIPG-11, and VUMC-HGG-09, were established through autopsy or resection material and characterized as described previously.^9^ Non-ATRT models JHH-DIPG-01, HSJD-DIPG-07, and SU-pcGBM-02 were kind gifts from Dr Raabe (Johns Hopkins Hospital, Baltimore, USA), Dr Montero Carcaboso (Hospital San Joan de Déu, Barcelona, Spain) and Dr Monje (Stanford University, Stanford, USA), respectively.

All cells were cultured as neurospheres as described previously.^9,11,12^ Cells were only used when confirmed mycoplasm negative and short-tandem repeat (STR) analysis was used to ensure cell line identities.

### Human mRNA expression datasets

Expression datasets that were used in this study to compare mRNA between patients were the following: Healthy brain (GSE: 11882)^13^, healthy cerebellum (GSE: 3526)^14^, healthy various (GSE: 7307, *Human body index project*; accession code: PRJNA98081), ATRT (GSE: 70678)^4^, glioblastoma (GSE: 7696)^15^, glioma (GSE: 16011)^16^, pediatric DIPG (GSE: 26576)^17^, pediatric glioma (GSE: 19578)^18^.

### Chemicals

The epigenetic compound library (#11076) was purchased from Cayman Chemical Company (Ann Arbor, Michigan, USA). Gemcitabine (CAS: 95058-81-4), doxorubicin (CAS: 23214-92-8), and staurosporine (62996-74-1) were purchased from MedChemExpress LLC (Monmouth Junction, New Jersey, USA). For *in vitro* studies, all chemicals were dissolved in DMSO and stored as 10mM stock concentration at −20 °C. For *in vivo* studies, both doxorubicin and gemcitabine where freshly dissolved in 0.9% saline before infusion.

### Compound screening and cell-viability assays

For compound screening assays, cells were plated in TSM-medium at a density of 5000 cells/well in cell-repellent 96-well F-bottom plates (#650971, Greiner Bio-one, Kremsmünster, Austria). Compounds were dispersed 24h after cell seeding, using a Tecan D300e picoliter dispenser (Tecan Group Ltd, Switzerland) and incubated at 37 °C and 5% CO2 for 96h. The number of viable cells was measured using CellTiter-Glo® 3D Luminescent Cell Viability Assay (#G9683, Promega, Madison, USA) according to the manufacturers protocol. The concordant luminescence was measured using a Tecan Infinite® 200 reader using iControl 1.10 software.

### Methylation profiling

DNA methylation arrays and copy number profiles were performed and analyzed using the Heidelberg classifier, as described previously.^4^

### RNA-sequencing

VUMC-ATRT-03, VUMC-DIPG-10, VUMC-DIPG-11, VUMC-HGG-09, JHH-DIPG-01, HSJD-DIPG-07, and SU-pcGBM-2 neurospheres were treated with 5nM gemcitabine or DMSO as control. After 24h, cells were collected and processed as described previously.^12^ Sequencing, performed on an Illumina Nextseq 500 sequencer, data processing in the R2 platform (R2.amc.nl), and statistical analysis were also assessed as previously described.^12,19^

### Western Blotting

Immunoblotting was performed as described previously.^20^ Protein isolation was conducted using RIPA lysis buffer supplemented with protease and phosphatase inhibitors. Membranes were incubated with mouse anti-SIRT1 (1:1000, #ab110304, Abcam, Cambridge, UK), rabbit anti-NF-kB p65 (D14E12) (1:1000, #8242, Cell Signaling Technology, Danvers, Massachusetts, USA), Rabbit anti-phospho-NF-kB p65 (Ser536, 93H1) (1:2000, #3033. Cell Signaling Technology), mouse anti-p53 (Clone DO-7) (1:1000, #M7001, DAKO, Agilent, Santa Clara, California, USA) and mouse anti-beta Actin (Clone C4) (1:10.000, MAB1501, Millipore, Burlington, Massachusetts, USA). Subsequently, membranes were incubated with secondary goat anti-mouse IRDye©600CV antibody (1:10.000, LI-COR©, Lincoln, NA, USA) and/or goat anti-rabbit IRDye©800CV antibody (1:10.000, LI-COR©). Signals were detected using a LI-COR© Odyssey fluorescent imager (model 9120, Surplus Solutions, LLC).

### Immunohistochemistry and immunofluorescent imaging

For immunohistochemistry, fresh tissues were fixed in 4% PFA for 48h, followed by paraffin embedding. Embedded tissues were cut into 4.5µm coronal sections and stained for SMARCB1 (1:100; 612111, BD Biosciences), Ki-67 (1:3000; ab15580, Abcam), glial fibrillary acidic protein (GFAP) (1:500; BT46-5002–04, BioTrend), S100 (1:1000; Z0311, DakoCytomation), human vimentin (1:4000; M0725, DakoCytomation), breast cancer resistance protein (BCRP) (1:1000; ab24115, Abcam), permeability glycoprotein (P-gp) (1:1000, #13978, Cell Signaling Technologies), CD31 (1:200, #PA5-16301, Thermo Fisher Scientific), p53 (1:500, Clone D07, M7001, DAKO), NF-kB (1:500, #D14E12, Cell Signaling technology), Sirtuin 1 (SIRT1) (1:100, #ab110304, Abcam, Cambridge, UK), glucose transporter 1 (GLUT1) (1:100–2000; 07-1401, Millipore) and hematoxylin for background contrast. Primary antibody detection was achieved through secondary peroxidase/DAB+ staining using the Dako REAL EnVision Detection System (Agilent, DAKO, #K5007).

For immunofluorescence, VUMC-ATRT-03 cells were cultured in TSM supplemented with FCS for 12h, to ensure adherence, in Greiner SCREENSTAR® 96-well plates (#655-866) specialized for fluorescent imaging. Cells were treated with gemcitabine for 24h, fixed and stained for GLI2 (1:100, ab223651, Abcam) as previously described.^12^ As secondary step Alexa Fluor™ 488 (goat anti-rabbit, 1:10,000, Invitrogen, #411-667) was incubated for 1h at RT. Imaging was performed using a Zeiss AxioObserver Z1 inverted microscope (Carl Zeiss, Oberkochen, Germany) using a 40x and 63x objective equipped with a Hamamatsu ORCA AG Black and White CCD camera (Hamamatsu Photonics, Shizuoka, Japan).

### Establishment of stable shRNA expressing cells

SIRT1 knockdown cells were established using the pLKO.1-shSIRT1.1, pLKO.1-shSIRT1.2, and pLKO.1-shSIRT1.3 (GE Healthcare, Chicago, USA) plasmids as described previously.^11^ pLKO.1-Scrambled plasmids (Addgene, #136035) were used as negative control. shRNA sequences are described in supplementary table S1.

### Establishment of stable TP53 knockout cells

VUMC-ATRT-01 cells were transduced (lentiviral) with a LentiCRISPR v2 plasmid (#52961, Addgene) that was cloned with a p53-sgRNA (Supplementary table S2) according to the applied protocol (#rev20140208, Zhang Lab). Using puromycin (2.0µg/mL) selection for 7 days, successfully transduced cells were selected. Knockout efficiency and location of the indel were confirmed through sanger sequencing, using TIDE analysis (tide.nki.nl) (Supplementary figure S1). Loss of p53 protein was assessed through western blotting.

### Animal ethics statement

All animal experiments were approved by local and governmental animal experimental committees and carried out according to national and institutional guidelines (national project permit: AVD114002017841) (protocol 0841-NCH17-01A1 and 0841-NCH20-13 animal welfare committee, Vrije Universiteit, Amsterdam).

### Xenograft modeling and *in vivo* efficacy studies

Both xenograft modeling and *in vivo* efficacy studies were performed in 4-week old female athymic nude mice (BALB/c outbred background, Envigo). Animals were provided food and water ad libitum for the entire duration of the experiments.

For modeling, mice were stereotactically injected with 500.000 patient-derived VUMC-ATRT-03 cells into the frontotemporal region of the left cerebral hemisphere (from bregma: X(−2), Y(0.5), Z(−3)mm), concordant with the original location of the tumor. Cells were injected in an injection volume of 5 μL at a flow rate of 2.5 μL/minute to minimize the neurological side effects of the procedure, as well as potential backflow of cells. Mice were euthanized upon developing severe neurological symptoms or >20% weight loss. Brains were fixed in 4% PFA for 48h for immunohistochemistry, or tumor cells were harvested and put in culture to establish VUMC-ATRT-03 mose passage cultures.

For *in vivo* efficacy studies, mice were stereotactically injected (as mentioned above) with 500.000 VUMC-ATRT-01 cells into the cerebellum (from lambda: X(−1), Y(−2.3), Z(−2.3)mm) or with 500.000 VUMC-ATRT-03 cells into the frontotemporal region of the left cerebral hemisphere, concordant with the original location in the patients. After eight days, mice were randomized into groups based on average body weight and received gemcitabine (200mg/kg), doxorubicin (20mg/kg) or 0.9% saline once per week for three weeks via I.P. injection. All animals developed tumors as confirmed by immunohistochemistry. Humane endpoints were set as either >20% weight loss or severe neurological symptoms.

### Statistics

mRNA expression between groups from RNA-sequencing data was assessed using the one-way analysis of variance (ANOVA). *In vitro* cell survival percentages were compared using the independent *t*-test (two-sided). *In vivo* survival differences between groups were tested using the log-rank (Mantel-Cox) test. The statistical analyzes were performed using GraphPad Prism (version 6). A p-value <0.05 was considered statistically significant.

## Results

### Establishment of a patient-derived ATRT culture and xenograft model

Tumor tissue was obtained through resection from an 11-year-old girl, suffering from a high-grade malignant, embryonal brain tumor. MRI imaging showed a large diffuse mass in the right fronto-temporal part of the cerebral hemisphere (Fig. 1A). Immunohistochemistry revealed complete loss of SMARCB1 in the nuclei of all tumor cells, corroborating the diagnosis ATRT (Fig. 1B). Genome-wide DNA methylation profiling confirmed the diagnosis and assigned the tumor to the SHH-subtype (supplementary figure S2). Fresh tumor tissue was put into culture (VUMC-ATRT-03) and maintained as neurospheres. Upon stable growth, VUMC-ATRT-03 cells were orthotopically injected into athymic nude mice. These mice developed tumors that, like the primary tumor, were characterized by a loss of SMARCB1 as shown by immunohistochemistry (Fig. 1C). Genome-wide DNA methylation profiling on VUMC-ATRT-03 cells harvested from mice confirmed the maintenance of the SHH-subtype within the model (Fig. S2). In addition, a panel of immunohistochemical stainings confirmed that high proliferation (Ki-67), multilineage differentiation (GFAP, S100, Vimentin) and loss of blood-brain-barrier integrity (GLUT1, BCRP, P-gp, CD31) were present in both original tumor tissue and mouse orthotopic xenografts of this neoplasm (Fig. 1D-E). Furthermore, chromosomal copy-number variations (CNV) as detected based on the genome-wide methylome analysis revealed that VUMC-ATRT-03 mouse passage model and original tumor tissue are largely overlapping, revealing that chromosomal alterations were maintained stable through modelling (Fig. 1F).

**Fig. 1.**
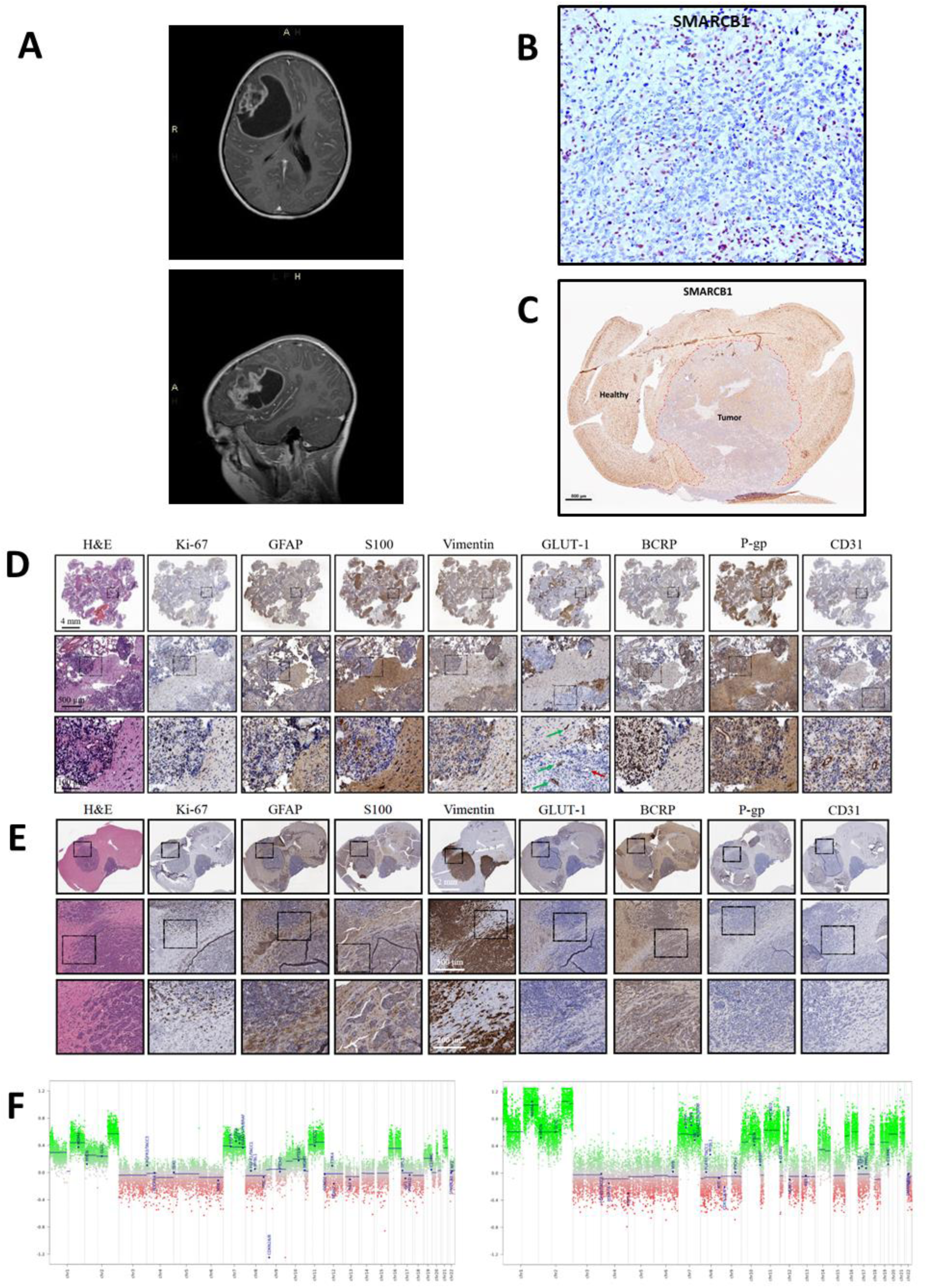
Establishment of a novel patient derived SHH-ATRT culture and xenograft model. (A) Diagnostic T2-weighted MRI of the patient from which VUMC-ATRT-03 was derived (upper panel: coronal plane / lower panel: sagittal plane). (B) Immunohistochemistry of patient-derived resection material depicting SMARCB1 expression in brown, revealing typical loss of nuclear SMARCB1 exclusively in tumor cells. The positive nuclei of non-neoplastic and microvascular cells serve as positive internal control. (C) Immunohistochemistry for SMARCB1 in a mouse brain carrying a VUMC-ATRT-03 xenograft confirming the absence of SMARCB1 in the nuclei of tumor cells. (D) Panel of immunohistochemical images depicting ATRT hallmarks in patient-derived resection material (high proliferation: KI-67; patches of multilineage differentiation: GFAP, S100, vimentin, keratin; loss of blood-brain-barrier integrity: GLUT-1, BCRP, P-gp, CD31). Loss of GLUT1 (indicated by the red arrow) indicates intratumor vascular malformations and loss of BBB integrity. (E) Panel of immunohistochemical images depicting ATRT hallmarks in a mouse brain carrying VUMC-ATRT-03 xenografts (high proliferation: KI-67; patches of multilineage differentiation: GFAP, S100, vimentin, keratin; loss of blood-brain-barrier integrity: GLUT-1, BCRP, P-gp, CD31). (F) Genomic copy number variation profile of VUMC-ATRT-03 patient-derived tumor tissue (left panel) and VUMC-ATRT-03 mouse passage cells (right panel).

### Gemcitabine is an effective ATRT-specific chemotherapeutic *in vitro*

We conducted an epigenetic compound-library screen (#11076, Cayman Chemical, USA) using the VUMC-ATRT-03 cells and a panel of ten other primary pediatric high-grade brain tumor neurosphere cultures. This screen indicated that only the ATRT model responded strongly to the clinically registered nucleoside analogue gemcitabine (Fig. 2A).^21^ To study if this sensitivity applied to more ATRT we tested the effect of gemcitabine on a panel of seven primary ATRT models and found all ATRT to be over a 10-fold more sensitive to gemcitabine compared to non-ATRT pediatric CNS-tumor models (Fig. 2B). Additionally, we observed that SHH-subtype ATRT models respond significantly more sensitive to gemcitabine treatment compared to MYC-subtype ATRT models.

**Fig. 2.**
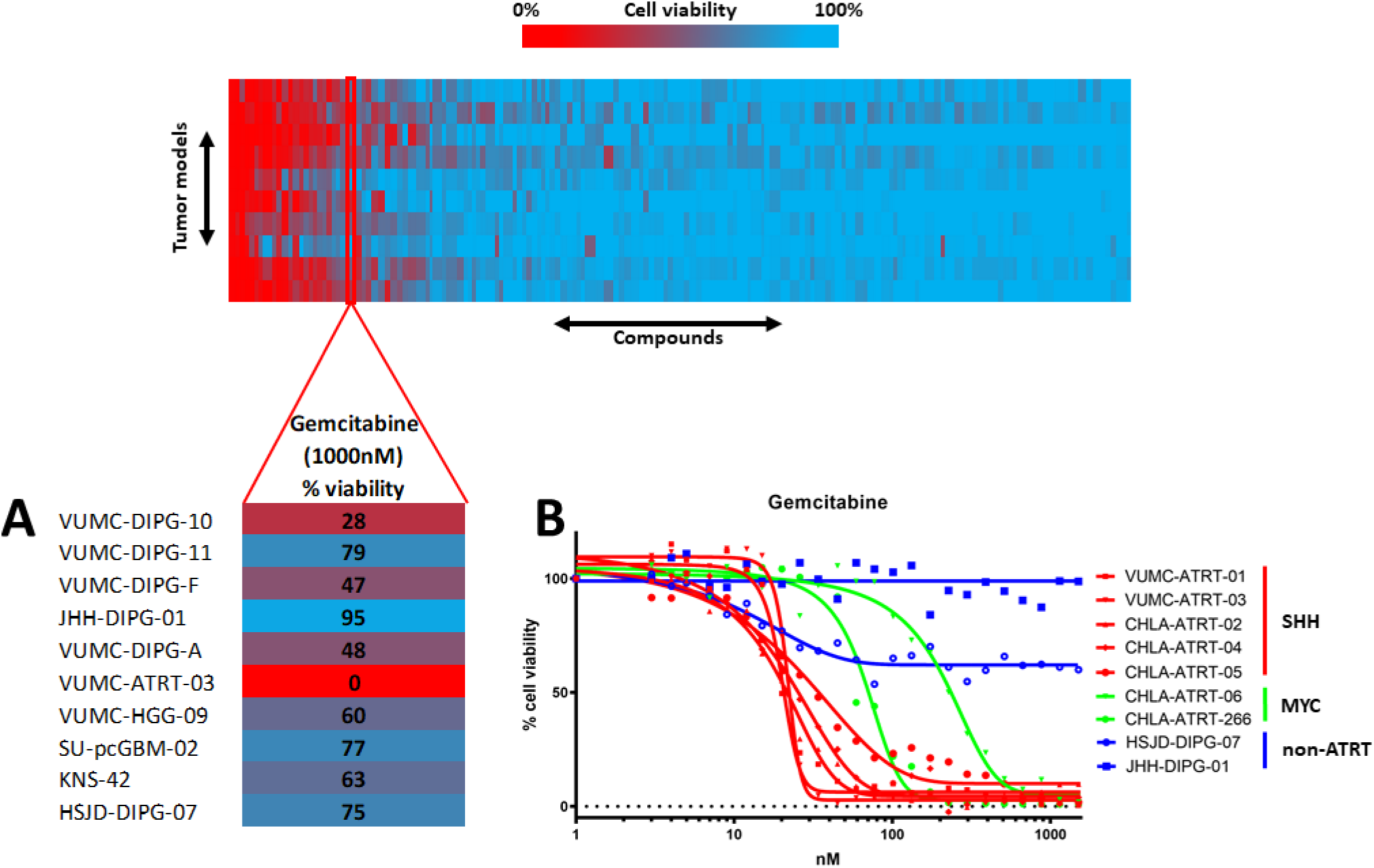
Epigenetic compound screening identifies gemcitabine as effective therapeutic in ATRT. (A) Compound library screen cell-viability readout in a panel of primary pediatric CNS tumor models. The gemcitabine treated cell-viability readout is highlighted in the lower panel, revealing that the only ATRT model from among the panel shows a complete loss of viability upon gemcitabine treatment (1000nM, 96h). (B) Cell-viability IC50 curves of gemcitabine treatment in a panel of seven primary ATRT culture models; two pediatric high-grade glioma cell cultures are used as controls. SHH-subtype ATRT models show a higher gemcitabine sensitivity compared to a MYC-subtype ATRT, while controls show no sensitivity.

### Gemcitabine treatment activates p53 signaling through depletion of ATRT-specific SIRT1 overexpression

To study the mechanisms underlying the ATRT-specific gemcitabine sensitivity we conducted RNA-sequencing after gemcitabine treatment. From analysis of this data we found that the nicotinamide adenosine dinucleotide-dependent histone deacetylase sirtuin-1 (*SIRT1*) is overexpressed in the VUMC-ATRT-03 model compared to non-ATRT pediatric brain-tumor models. Furthermore, *SIRT1* RNA expression was found to be significantly elevated solely in the ATRT model upon gemcitabine treatment (Fig. 3A). To investigate if SIRT1 overexpression applies to all ATRT, we looked at publicly available gene-expression datasets (R2 platform) and compared SIRT1 expression in 49 ATRT samples to healthy brain and cerebellum and to other high-grade brain tumor tissues. We found that SIRT1 is significantly overexpressed in ATRT compared to healthy brain tissues and other brain tumor samples (Fig. 3B). In contrast to RNA-expression, western blot analysis revealed that gemcitabine treatment depletes SIRT1 protein expression in primary ATRT cell cultures *in vitro* (Fig. 3C), which may indicate a transcriptional feedback loop upon SIRT1 depletion. In parallel we found that gemcitabine treatment upregulates mRNA expression of both Nuclear Factor-kappa-B (NF-kB) subunits, NFKB1 and NFKB2, solely in the ATRT model (Fig. 3D). Furthermore, RNA-expression data revealed that gemcitabine upregulates transcription of the full NF-kB signaling pathway (Fig. 3E). Using western blotting we confirmed that gemcitabine treatment activates the NF-kB signaling pathway by increasing the phosphorylation of the p65 NF-kB subunit in our primary ATRT models (Fig. 3F). Since SIRT1 and NF-kB are both modulators of p53, and the majority of ATRT do not harbor a p53 mutation, we hypothesized that p53 may play a role in ATRT specific gemcitabine sensitivity.^22–25^ Therefore, we assessed p53 protein expression in five primary ATRT models, before and after gemcitabine treatment, and found that all ATRT models show elevated p53 protein expression upon gemcitabine treatment (Fig. 3G). Furthermore, RNA-expression data revealed that gemcitabine treatment activates the overall (KEGG defined) p53 signaling machinery, as well as p53 transcriptional targets, in the VUMC-ATRT-03 model (Fig. 3H and Supplementary figure S3). Additionally, since ATRT-patients often receive NF-kB suppressing corticosteroids, we tested if dexamethasone would suppress gemcitabine effectivity in our ATRT models. No anti-synergism was found between dexamethasone and gemcitabine at clinically relevant doses (Supplementary figure S4).

**Fig. 3.**
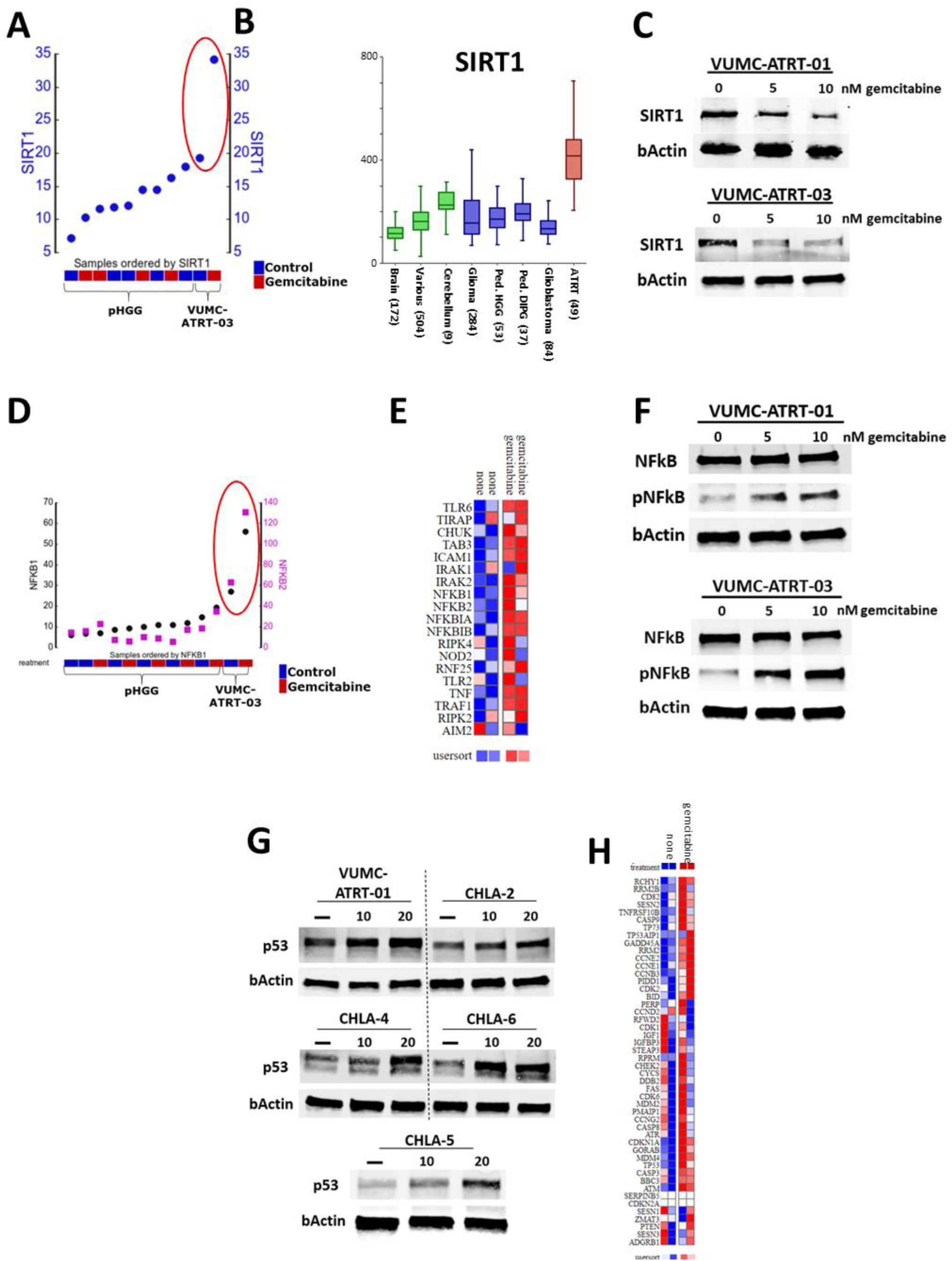
ATRT specific SIRT1 upregulation suppresses NF-kB and p53 activity, which can be reversed through gemcitabine treatment. (A) Untreated (blue) vs. gemcitabine (red) treated mRNA expression of SIRT1 in VUMC-ATRT-03 and a panel of pediatric HGG models. (B) mRNA expression levels of SIRT1 in normal brain, cerebellum and various tissues (in green, GSE: 11882, GSE: 3526, GSE: 7307) compared to adult and pediatric CNS tumor tissues (in blue, GSE: 7696, GSE: 16011, GSE: 26576, GSE: 19578) and ATRT tissues (in red, GSE: 70678). (C) Western blot analysis depicting SIRT1 expression in VUMC-ATRT-01 and VUMC-ATRT-03 cells after gemcitabine treatment (0, 5, and 10nM) for 24h. (D) Untreated (blue) vs. gemcitabine (red) treated mRNA expression of NFKB1 (black) and NFKB2 (magenta) in VUMC-ATRT-03 and a panel of pediatric HGG models. (E) Heatmap representation illustrating mRNA expression of the KEGG NF-kB gene set from RNA-sequencing data showing two non-treated versus gemcitabine treated ATRT cultures (PAGE FDR *P*<0.05). (F) Western blot analysis depicting p65-NFkB and phospho-p65-NFkB (pNFkB) expression in VUMC-ATRT-01 and VUMC-ATRT-03 cells after gemcitabine treatment (0, 5, and 10nM) for 24h. (G) Western blot analysis depicting p53 expression in VUMC-ATRT-01, VUMC-ATRT-03, CHLA-ATRT-02, CHLA-ATRT-04, CHLA-ATRT-05, and CHLA-ATRT-06 cells after gemcitabine treatment (0, 10, and 20nM) for 24h. (H) Heatmap representation illustrating mRNA expression of the KEGG p53 signaling gene set from RNA-sequencing data showing two non-treated versus gemcitabine treated ATRT cultures (PAGE FDR *P*<0.05).

### Gemcitabine-induced SIRT1 depletion causes p53-mediated cell death in ATRT

To study the role of SIRT1 and p53 in ATRT cells we engineered VUMC-ATRT-01 and VUMC-ATRT-03 cells in which SIRT1 is downregulated by stable shRNA expression (Fig. 4A) and used CRISPR/Cas9 to make stable p53 knockouts of these models (Fig. 4C). The *SIRT1* knockdown in ATRT cells did not show loss of cell-viability in culture, but when treated with gemcitabine we observed an average 10-fold increased sensitivity compared to wild-type ATRT (Fig. 4B). Adversely, *p53* knockout in ATRT cells caused a complete loss of sensitivity to gemcitabine (Fig. 4D), demonstrating the pivotal role of p53 in gemcitabine toxicity in ATRT (Fig. 4E).

**Fig. 4.**
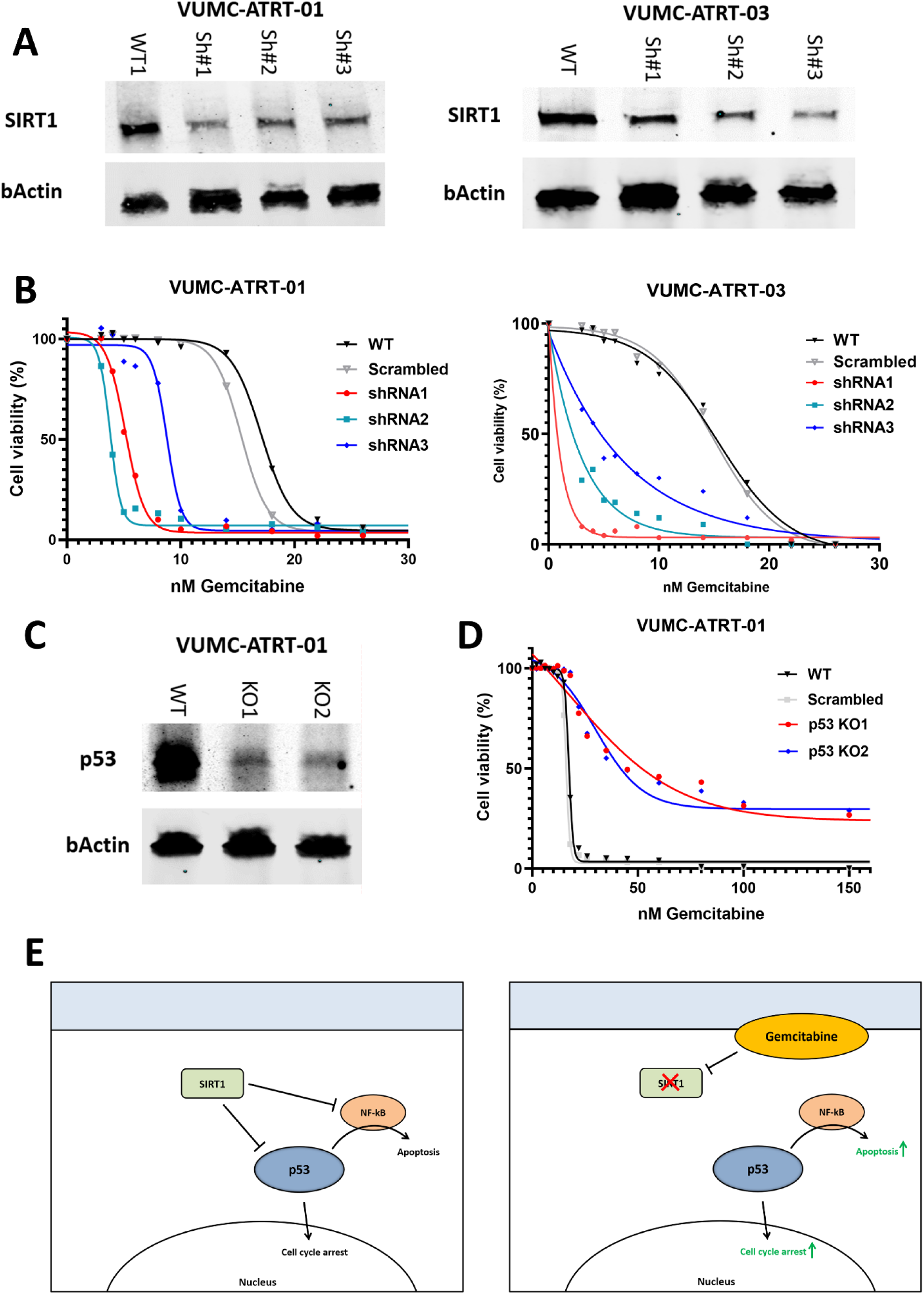
Gemcitabine treatment induces p53 mediated cell-death, through SIRT1 depletion in ATRT. (A) Western blot analysis depicting SIRT1 expression in VUMC-ATRT-01 and VUMC-ATRT-03 cells transduced with a variety of SIRT1 shRNA (Sh#1, 2, and 3) or a control construct (WT). (B) IC50 viability curves of VUMC-ATRT-01 and VUMC-ATRT-03 cells transduced with control (scrambled) or one of three SIRT1 shRNA treated with different gemcitabine concentrations for 96h. (C) Western blot analysis depicting p53 expression in VUMC-ATRT-01 wildtype (WT) and p53 knockout cells (KO1 and KO2), as established through CRISPR/Cas9. (D) IC50 viability curves of CRISPR/Cas9 modified VUMC-ATRT-01 WT, scrambled, and p53 knockout cells, treated with different gemcitabine concentration for 96h. (E) Illustration of the proposed mechanism through which gemcitabine causes tumor toxicity in ATRT.

### Gemcitabine blocks SHH activity through loss of GLI2 nuclear localization

As described above, we observed differential gemcitabine sensitivity between the SHH-ATRT and MYC-ATRT models. Therefore, we investigated differential mRNA expression between SHH-ATRT and non-SHH-ATRT in a publicly available expression dataset consisting of 49 ATRT samples (Fig. 5A). As anticipated, we identified SIRT1 as being significantly upregulated in SHH-compared to non-SHH-ATRT (Fig. 5B). Besides its role as a p53 suppressor, SIRT1 is also known to be an activator of GLI2, the hallmark protein for SHH signaling activity, through GLI2 deacetylation which allows for nuclear translocation.^26^ As such, we hypothesized that gemcitabine-induced loss of SIRT1 would inhibit GLI2 deacetylation, thus preventing nuclear localization of GLI2. Therefore, we performed GLI2 immunofluorescence staining in VUMC-ATRT-03 cells, before and after gemcitabine treatment, and found that this treatment causes a significant loss of GLI2 nuclear localization, indicating lost SHH-signaling activity (Fig. 5C). Furthermore, parametric analysis of gene set enrichment (PAGE) revealed negative regulation of expression of genes involved in SHH signaling in our cultures upon treatment with gemcitabine (Supplementary table S3). These results indicate that gemcitabine-induced loss of SIRT1 acts as a double-edged sword in SHH-ATRT through both activation of p53 and inhibition of SHH signaling activity (Fig. 5D).

**Fig. 5.**
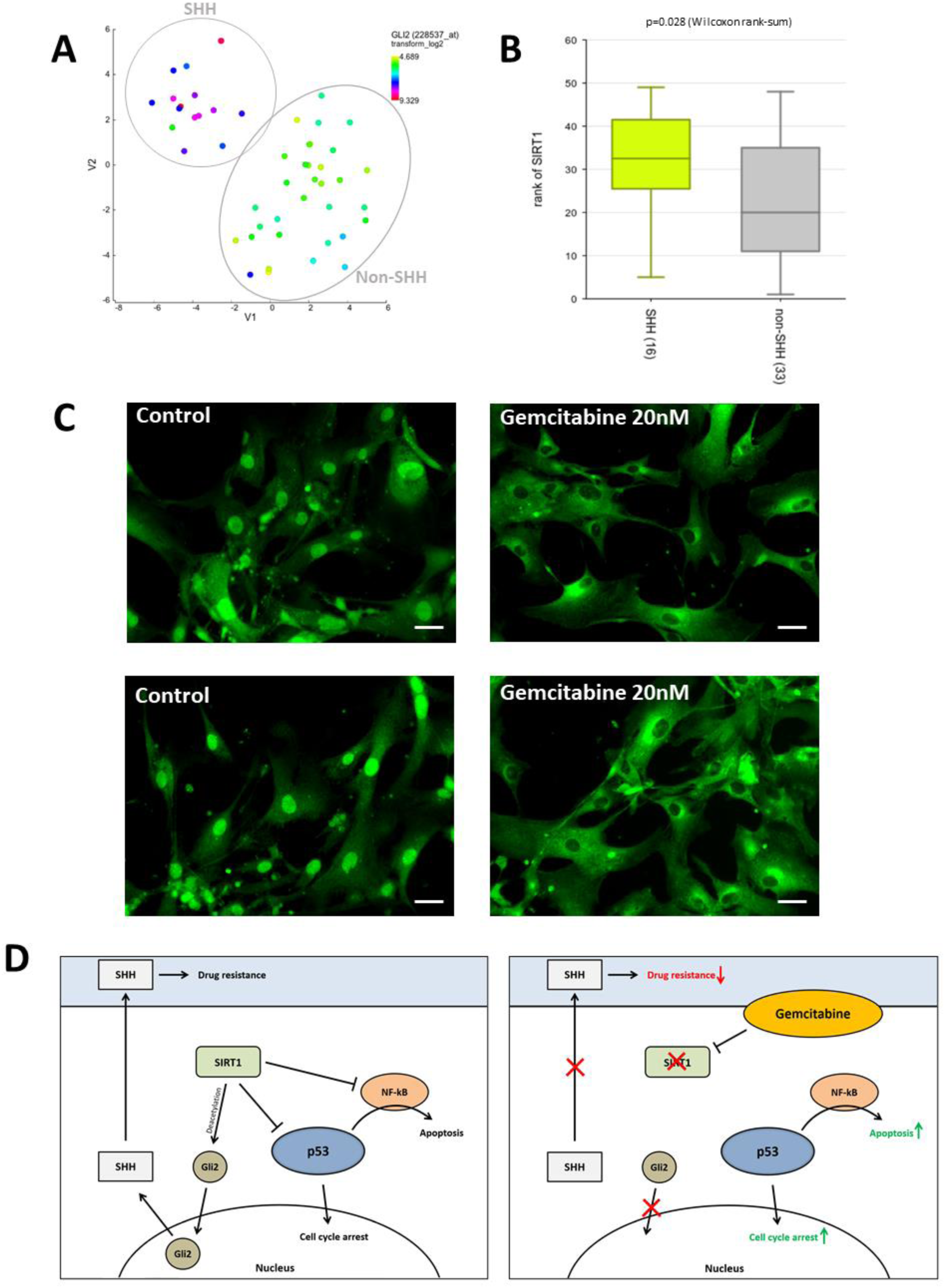
Gemcitabine deactivates SHH signaling in SHH-ATRTs through loss of SIRT1, increasing SHH-ATRT specific drug-sensitivity. (A) T-distributed Stochastic Neighbor Embedding (t-SNE) clustering of 49 individual ATRT patient tumor samples (dataset GSE: 70678) based on overall mRNA expression profiles (perplexity: 12). SHH-subgroup is shown as a distinct group from other ATRT samples in upper-left corner as confirmed by overall high GLI2 expression (blue to red) versus low Gli2 expression in the non-SHH ATRT (yellow to green). (B) mRNA expression levels of SIRT1 in SHH ATRT versus non-SHH ATRT. (C) Immunofluorescent stainings of GLI2 in VUMC-ATRT-03 cells treated with 20nM gemcitabine for 24h shows that loss of nuclear localization of GLI2. Scale bar indicates 20µm. (D) Illustration of the proposed mechanism through which gemcitabine causes tumor toxicity in ATRT, including the mechanisms that causes extra sensitivity in SHH-subgroup ATRT.

### Gemcitabine treatment prolongs survival in ATRT xenograft models

To evaluate the therapeutic efficacy of gemcitabine *in vivo*, we tested this compound in athymic nude mice carrying patient-derived ATRT xenografts (VUMC-ATRT-03). Treatment (14 days) started eight days after tumor transplantation, at which mice were stratified into two equivalent groups (control and gemcitabine) based on average trend in body weight. No treatment-related toxicities were observed, and all mice developed ATRT, as determined by *post-mortem* immunohistochemistry. We observed over 30% prolonged survival (p=0.003) in the gemcitabine group compared to control mice, based on >20% weight-loss or severe neurological symptoms as exclusion criteria (Fig. 6A-B). Besides the frontotemporal ATRT xenograft model VUMC-ATRT-03, we also tested therapeutic efficacy of gemcitabine in an orthotopic xenograft of a cerebellar ATRT model (VUMC-ATRT-01). To this second *in vivo* trial we added a doxorubicin group as positive treatment control. Here we found a similar 30% prolonged survival (p=0.0008) of gemcitabine treated mice compared to both control and doxorubicin treated mice (Fig. 6C-D). Additionally, in both trials we found one long-term survivor among gemcitabine treated mice, which were sacrificed upon the determined endpoint of each trial (day 120 and 180 respectively). To evaluate the hypothesized underlying mechanisms of gemcitabine efficacy, we collected brains from five pre-selected mice at the end of their treatment cycles on which we conducted immunohistochemistry. In line with our *in vitro* data, we found that gemcitabine treatment activates p53 (Fig. 6E), while causing a loss of SIRT1, concordant with our previous findings (Fig. 6F). Immunohistochemistry also confirmed that all mice, including both long-term survivors, developed ATRT.

**Fig. 6.**
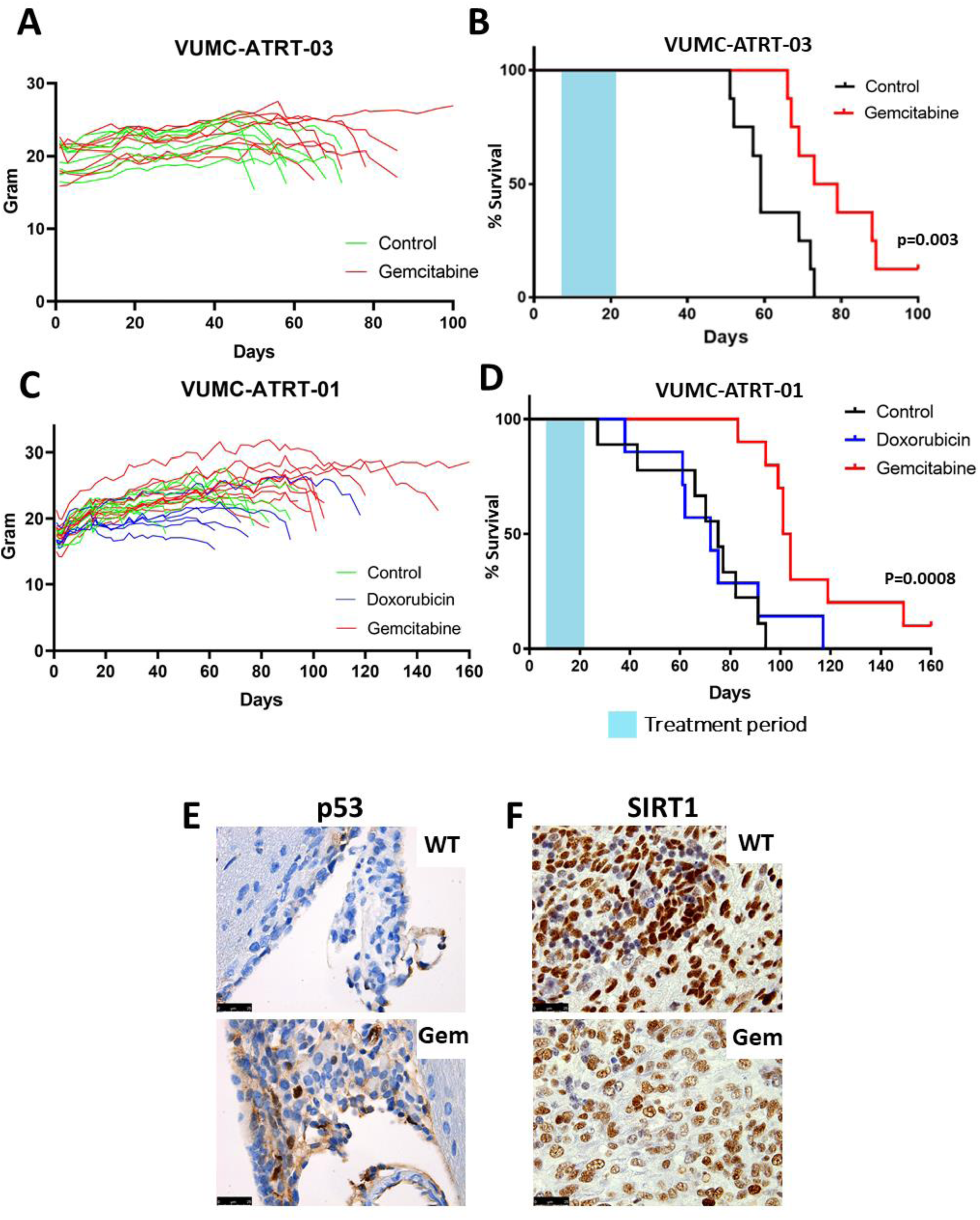
Gemcitabine treatment reveals prolonged survival in two SHH-ATRT xenograft models. (A) Weight development of VUMC-ATRT-03 orthotopic xenograft bearing mice over time (control in green, gemcitabine treated in red). (B) Survival analysis of VUMC-ATRT-03 orthotopic xenograft bearing mice treated with vehicle (black line, n=8) and gemcitabine (red line, n=8). The gemcitabine treated group shows significant survival benefit over vehicle treated mice (p=0.003, log-rank test). (C) Weight development of VUMC-ATRT-01 orthotopic xenograft bearing mice over time (control in green, doxorubicin treated in blue, gemcitabine treated in red). (D) Survival analysis of VUMC-ATRT-01 orthotopic xenograft bearing mice treated with vehicle (black line, n=9), doxorubicin (blue line, n=7) and gemcitabine (red line, n=10). The gemcitabine treated group shows significant benefit over vehicle and doxorubicin treated mice (p=0.0008, log-rank test). (E) p53 immunohistochemical staining (in brown) in VUM-ATRT-01 xenograft patches, isolated at final day of treatment. Gemcitabine treated mice show a higher ratio of p53 positive nuclei in their tumors compared to vehicle treated animals. (F) SIRT1 immunohistochemical staining (in brown) in VUM-ATRT-01 xenograft patches, isolated at final day of treatment. Gemcitabine treated mice show lower SIRT1 expression compared to vehicle treated animals.

## Discussion

In the current study we show that gemcitabine selectively acts as a potent chemotherapeutic agent in ATRT compared to non-ATRT *in vitro* and *in vivo* pediatric brain tumor models. From expression data we observed that this sensitivity correlates to the overexpression of *SIRT1* mRNA, which is thought to act as a promotor of tumorigenesis and drug resistance by hypermethylating the promotor regions of several tumor suppressor genes. ^27–30^.

Gemcitabine is a clinically well-known chemotherapeutic that was approved for medical use in 1995 and was even added to the World Health Organization’s List of Essential Medicines.^31^ Several subsequent studies investigated brain penetration of gemcitabine and found that gemcitabine does cross the blood-brain-barrier (BBB), albeit to a limited extend. However, the gemcitabine concentrations found in glioblastoma biopsies, taken from patients treated with a clinically relevant dose of 1000mg/m^2^, exceeded the lethal gemcitabine dose for our SHH-ATRT *in vitro* models.^32^ In pediatric patients, weekly gemcitabine doses up to 2100 mg/m^2^ are well tolerated, suggesting that relevant gemcitabine brain concentrations in these patients should be feasible.^33^ Additionally, in a previous study we found that ATRT share significant physiological BBB deficiencies and that even non-BBB penetrable drugs can penetrate ATRT in xenograft models.^10^ As such, we do expect that gemcitabine penetrates the brain and can be used for potential clinical treatment of ATRT.

The tumor suppressor p53 is considered the main inducer of cell-cycle arrest and apoptosis, and the vast majority of cancers harbor either a p53 mutation or have lost the ability to induce p53. In this study we show that gemcitabine mainly causes its cytotoxic effect in ATRT cells through activation of p53. These results indicate that gemcitabine would only be effective in p53 wild-type tumors, which underlines the clinical relevance of this study, since ATRT rarely carry p53 mutations.^8^ Furthermore, we show that the p53 inhibitor SIRT1 is overexpressed in ATRT compared to other CNS tumors, which might indicate why p53 is often suppressed in ATRT, contributing to its highly malignant behavior.

In correlation with p53, we found that NF-kB is activated upon gemcitabine treatment in our ATRT models and we suggest that this activation works synergistically with p53 to induce apoptosis. For decades there has been discussion on the pro-apoptotic versus the anti-apoptotic roles of NF-kB signaling. Despite contradictory studies that have shown both anti- and pro-apoptotic functioning of NF-kB activity, even within the same cell lines or cancers, today there is an overall consensus that NF-kB is a context-dependent apoptosis regulator.^24,34,35^ As a modulator of apoptosis, excessive NF-kB activation has been described to contribute to p53-regulated apoptosis in several cancers, corresponding to our observations in this study.^36^

Since some other aggressive tumors like acute myeloid leukemia, colon cancer and pancreatic cancer also overexpress SIRT1, gemcitabine treatment may also be applicable to those tumor types. This is confirmed by the observation that a direct link between SIRT1 expression and gemcitabine treatment efficacy has been found in pancreatic cancers.^37–39^ These thorough and exclusive studies show that pancreatic ductal adenocarcinoma rely on high SIRT1 expression for their proliferation and migration capacities and that inhibition of SIRT1 directly sensitizes these tumors to gemcitabine treatment.

Interestingly, like SHH-ATRT, pancreatic cancers often are characterized by highly active hedgehog signaling.^39,40^ Several recent studies show that gemcitabine resistance in pancreatic tumors is the result of reactivation of SHH signaling, which induces stemcellness and embryonal development.^41,42^ These results support our findings that show how loss of SHH-signaling in SHH-subtype ATRT causes strong gemcitabine sensitivity.

In this study, systemic treatment of murine ATRT orthotopic xenograft models with gemcitabine prolonged survival significantly. As such, in the second *in vivo* trial we compared the efficacy of monotherapy gemcitabine with that of doxorubicin, a currently used chemotherapeutic agent in ATRT treatment regimens.^2,43^ Our results show that gemcitabine monotherapy causes a significantly stronger antitumoral effect in ATRT than doxorubicin, without causing any adverse effects *in vivo*, which alludes to the potential of gemcitabine as part of multimodal therapy in ATRT.

In conclusion, we demonstrate that ATRT are highly sensitive to the clinically registered chemotherapeutic gemcitabine. We found that gemcitabine treatment hampers ATRT specific SIRT1 overexpression, causing p53 activation and cell death in these tumors. Additionally, we demonstrate that gemcitabine sensitivity is more pronounced in SHH-subtype ATRT, supposedly because of the GLI2 activating role of SIRT1, which is diminished upon gemcitabine treatment. Finally, we demonstrate that gemcitabine significantly prolongs survival of mice with orthotopic ATRT xenografts compared to both control and conventional doxorubicin treatment. Overall, gemcitabine treatment may offer an easily translatable opportunity to improve outcomes for ATRT patients, for whom effective treatment option are still scarce.

## Notes

**Conflict of interest:** None of the authors declare any conflicts of interest.

### Competing Interest Statement

The authors have declared no competing interest.

## References

1. Ginn KF, Gajjar A. Atypical teratoid rhabdoid tumor: current therapy and future directions. Front Oncol. 2012; 2:114.

2. Slavc I, Chocholous M, Leiss U, et al. Atypical teratoid rhabdoid tumor: improved long-term survival with an intensive multimodal therapy and delayed radiotherapy. The Medical University of Vienna Experience 1992-2012. Cancer Med. 2014; 3(1):91–100.

3. Hasselblatt M, Gesk S, Oyen F, et al. Nonsense mutation and inactivation of SMARCA4 (BRG1) in an atypical teratoid/rhabdoid tumor showing retained SMARCB1 (INI1) expression. Am J Surg Pathol. 2011; 35(6):933–935.

4. Johann PD, Erkek S, Zapatka M, et al. Atypical Teratoid/Rhabdoid Tumors Are Comprised of Three Epigenetic Subgroups with Distinct Enhancer Landscapes. Cancer Cell. 2016; 29(3):379–393.

5. Torchia J, Golbourn B, Feng S, et al. Integrated (epi)-Genomic Analyses Identify Subgroup-Specific Therapeutic Targets in CNS Rhabdoid Tumors. Cancer Cell. 2016; 30(6):891–908.

6. Ho B, Johann PD, Grabovska Y, et al. Molecular subgrouping of atypical teratoid/rhabdoid tumors-a reinvestigation and current consensus. Neuro Oncol. 2020; 22(5):613–624.

7. Abro B, Kaushal M, Chen L, et al. Tumor mutation burden, DNA mismatch repair status and checkpoint immunotherapy markers in primary and relapsed malignant rhabdoid tumors. Pathol Res Pract. 2019; 215(6):152395.

8. Gröbner SN, Worst BC, Weischenfeldt J, et al. The landscape of genomic alterations across childhood cancers. Nature. 2018; 555(7696):321-327.

9. Meel MH, Sewing ACP, Waranecki P, et al. Culture methods of diffuse intrinsic pontine glioma cells determine response to targeted therapies. Exp Cell Res. 2017; 360(2):397–403.

10. Meel MH, Guillén Navarro M, de Gooijer MC, et al. MEK/MELK inhibition and blood-brain barrier deficiencies in atypical teratoid/rhabdoid tumors. Neuro Oncol. 2020; 22(1):58–69.

11. Meel MH, Metselaar DS, Waranecki P, Kaspers GJL, Hulleman E. An efficient method for the transduction of primary pediatric glioma neurospheres. MethodsX. 2018; 5:173–183.

12. Metselaar DS, Meel MH, Benedict B, et al. Celastrol-induced degradation of FANCD2 sensitizes pediatric high-grade gliomas to the DNA-crosslinking agent carboplatin. EBioMedicine. 2019; 50:81–92.

13. Berchtold NC, Cribbs DH, Coleman PD, et al. Gene expression changes in the course of normal brain aging are sexually dimorphic. Proc Natl Acad Sci U S A. 2008; 105(40):15605–15610.

14. Roth RB, Hevezi P, Lee J, et al. Gene expression analyses reveal molecular relationships among 20 regions of the human CNS. Neurogenetics. 2006; 7(2):67–80.

15. Murat A, Migliavacca E, Gorlia T, et al. Stem cell-related “self-renewal” signature and high epidermal growth factor receptor expression associated with resistance to concomitant chemoradiotherapy in glioblastoma. J Clin Oncol. 2008; 26(18):3015–3024.

16. Gravendeel LA, Kouwenhoven MC, Gevaert O, et al. Intrinsic gene expression profiles of gliomas are a better predictor of survival than histology. Cancer Res. 2009; 69(23):9065–9072.

17. Paugh BS, Broniscer A, Qu C, et al. Genome-wide analyses identify recurrent amplifications of receptor tyrosine kinases and cell-cycle regulatory genes in diffuse intrinsic pontine glioma. J Clin Oncol. 2011; 29(30):3999–4006.

18. Paugh BS, Qu C, Jones C, et al. Integrated molecular genetic profiling of pediatric high-grade gliomas reveals key differences with the adult disease. J Clin Oncol. 2010; 28(18):3061–3068.

19. Meel MH, de Gooijer MC, Metselaar DS, et al. Combined Therapy of AXL and HDAC Inhibition Reverses Mesenchymal Transition in Diffuse Intrinsic Pontine Glioma. Clin Cancer Res. 2020; 26(13):3319–3332.

20. Meel MH, de Gooijer MC, Guillén Navarro M, et al. MELK Inhibition in Diffuse Intrinsic Pontine Glioma. Clin Cancer Res. 2018; 24(22):5645–5657.

21. de Sousa Cavalcante L, Monteiro G. Gemcitabine: metabolism and molecular mechanisms of action, sensitivity and chemoresistance in pancreatic cancer. Eur J Pharmacol. 2014; 741:8–16.

22. Vaziri H, Dessain SK, Ng Eaton E, et al. hSIR2(SIRT1) functions as an NAD-dependent p53 deacetylase. Cell. 2001; 107(2):149–159.

23. Webster GA, Perkins ND. Transcriptional cross talk between NF-kappaB and p53. Mol Cell Biol. 1999; 19(5):3485–3495.

24. Lin B, Williams-Skipp C, Tao Y, et al. NF-kappaB functions as both a proapoptotic and antiapoptotic regulatory factor within a single cell type. Cell Death Differ. 1999; 6(6):570–582.

25. Lin Z, Fang D. The Roles of SIRT1 in Cancer. Genes Cancer. 2013; 4(3-4):97–104.

26. Xie Y, Liu J, Jiang H, et al. Proteasome inhibitor induced SIRT1 deacetylates GLI2 to enhance hedgehog signaling activity and drug resistance in multiple myeloma. Oncogene. 2020; 39(4):922–934.

27. Sasca D, Hähnel PS, Szybinski J, et al. SIRT1 prevents genotoxic stress-induced p53 activation in acute myeloid leukemia. Blood. 2014; 124(1):121–133.

28. Jiang K, Lyu L, Shen Z, et al. Overexpression of SIRT1 is a poor prognostic factor for advanced colorectal cancer. Chin Med J (Engl). 2014; 127(11):2021–2024.

29. Stenzinger A, Endris V, Klauschen F, et al. High SIRT1 expression is a negative prognosticator in pancreatic ductal adenocarcinoma. BMC Cancer. 2013; 13:450.

30. Wang Z, Chen W. Emerging Roles of SIRT1 in Cancer Drug Resistance. Genes Cancer. 2013; 4(3-4):82–90.

31. World Health O. World Health Organization model list of essential medicines: 21st list 2019. Geneva: World Health Organization; 2019 2019.

32. Sigmond J, Honeywell RJ, Postma TJ, et al. Gemcitabine uptake in glioblastoma multiforme: potential as a radiosensitizer. Annals of Oncology. 2009; 20(1):182–187.

33. Reid JM, Qu W, Safgren SL, et al. Phase I trial and pharmacokinetics of gemcitabine in children with advanced solid tumors. J Clin Oncol. 2004; 22(12):2445–2451.

34. Kaltschmidt B, Kaltschmidt C, Hofmann TG, Hehner SP, Dröge W, Schmitz ML. The pro- or anti-apoptotic function of NF-kappaB is determined by the nature of the apoptotic stimulus. Eur J Biochem. 2000; 267(12):3828–3835.

35. Radhakrishnan SK, Kamalakaran S. Pro-apoptotic role of NF-kappaB: implications for cancer therapy. Biochim Biophys Acta. 2006; 1766(1):53–62.

36. Ryan KM, Ernst MK, Rice NR, Vousden KH. Role of NF-kappaB in p53-mediated programmed cell death. Nature. 2000; 404(6780):892-897.

37. Gong DJ, Zhang JM, Yu M, Zhuang B, Guo QQ. Inhibition of SIRT1 combined with gemcitabine therapy for pancreatic carcinoma. Clin Interv Aging. 2013; 8:889–897.

38. Zhao G, Cui J, Zhang JG, et al. SIRT1 RNAi knockdown induces apoptosis and senescence, inhibits invasion and enhances chemosensitivity in pancreatic cancer cells. Gene Ther. 2011; 18(9):920–928.

39. Oon CE, Strell C, Yeong KY, Östman A, Prakash J. SIRT1 inhibition in pancreatic cancer models: contrasting effects in vitro and in vivo. Eur J Pharmacol. 2015; 757:59–67.

40. Bai Y, Bai Y, Dong J, et al. Hedgehog Signaling in Pancreatic Fibrosis and Cancer. Medicine (Baltimore). 2016; 95(10):e2996.

41. Jia Y, Gu D, Wan J, et al. The role of GLI-SOX2 signaling axis for gemcitabine resistance in pancreatic cancer. Oncogene. 2019; 38(10):1764–1777.

42. Jia Y, Xie J. Promising molecular mechanisms responsible for gemcitabine resistance in cancer. Genes Dis. 2015; 2(4):299–306.

43. Chi SN, Zimmerman MA, Yao X, et al. Intensive multimodality treatment for children with newly diagnosed CNS atypical teratoid rhabdoid tumor. J Clin Oncol. 2009; 27(3):385–389.

