## Supplemental figures for "Gemcitabine therapeutically disrupts essential SIRT1-mediated p53 repression in Atypical Teratoid/Rhabdoid Tumors"

**Supplementary figures**

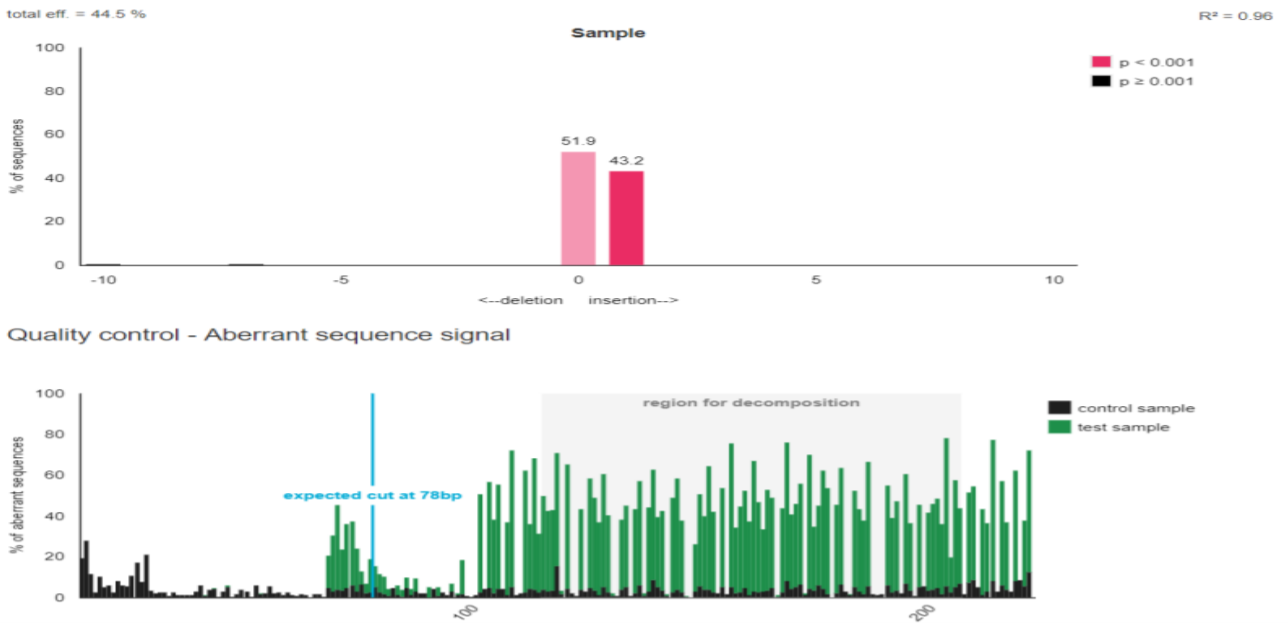

***Supplementary figure S1:* TP53 indel efficiency in VUMC-ATRT-03 cells upon Cas9 and TP53 sgRNA transduction.**
Upper panel: decomposition yielding the spectrum of indels and their frequencies (before blasticidin selection) as analyzed by TIDE. Lower panel: visualization of aberrant sequence signal in control (black) and treated sample (green). The region used for decomposition is indicated with a gray bar; the expected break site with a vertical blue line.

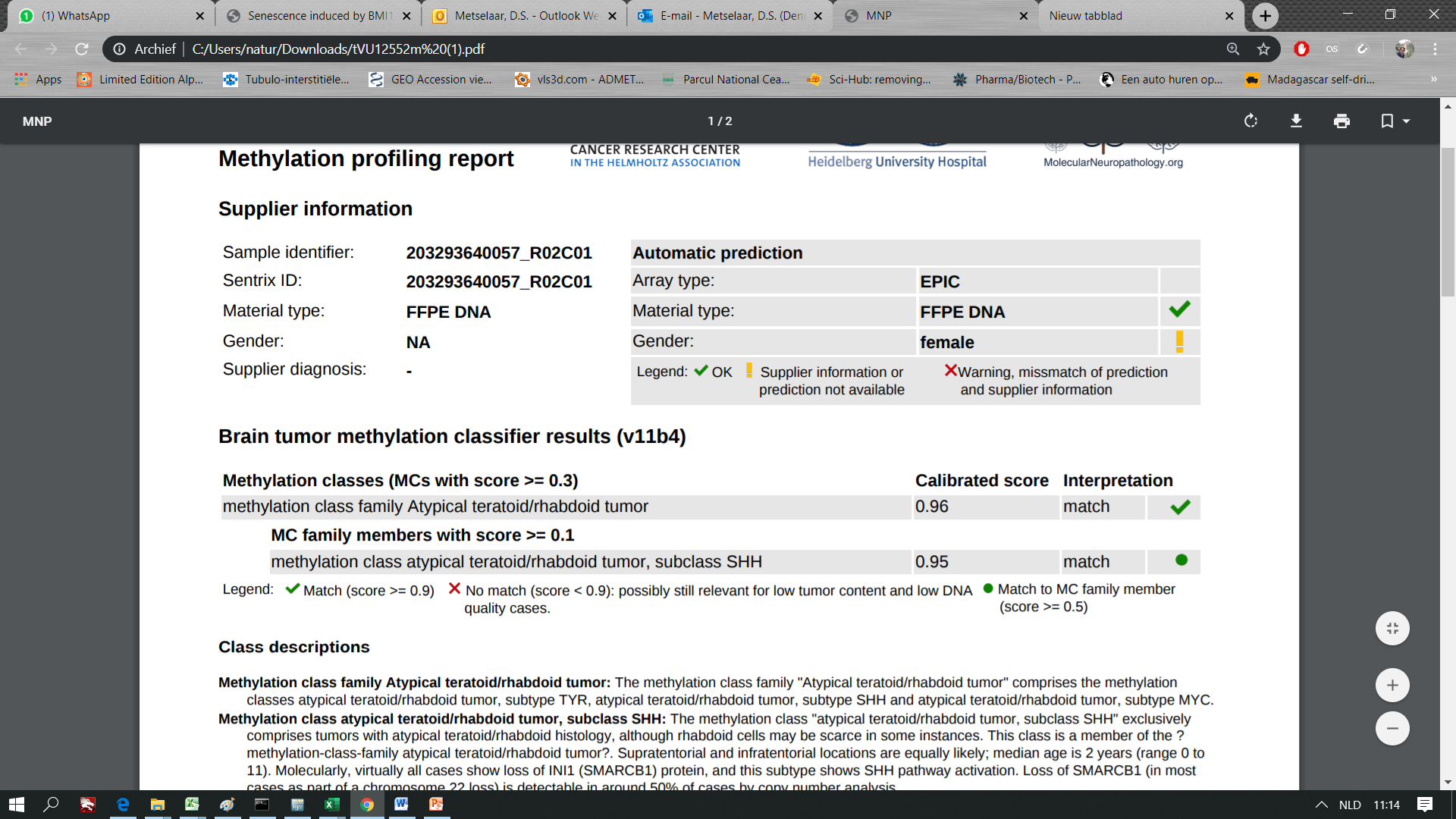

***Supplementary figure S2:*** Methylome profiling analysis of the VUMC-ATRT-03 cells and matching of this profile with the Heidelberg Brain Tumors Classifier^44^ (PMID 29539639; [www.molecularneuropathology.org](http://www.molecularneuropathology.org)) confirms that these cells belong to the SHH subgroup of ATRT.

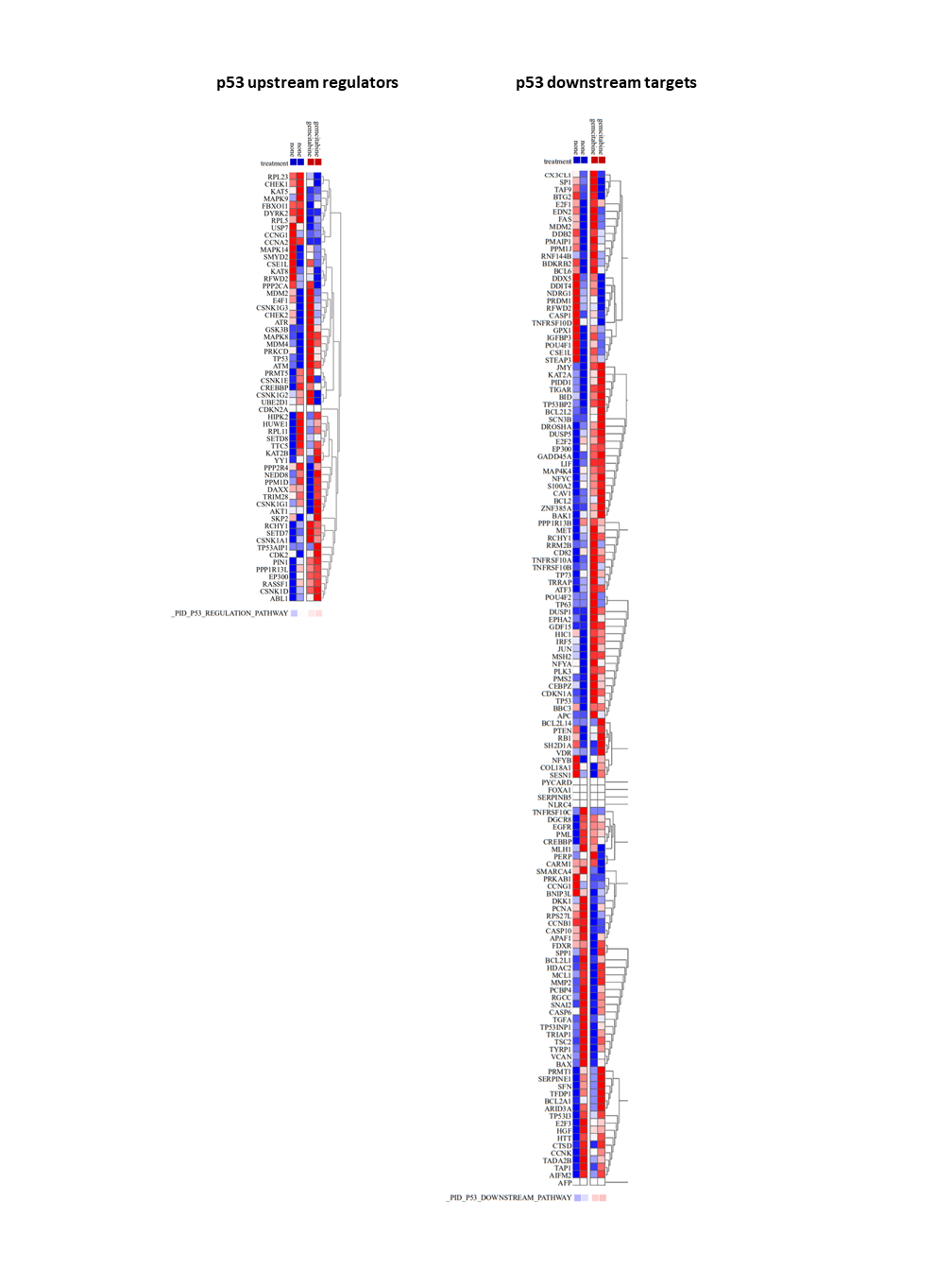

***Supplementary figure S3:*** Heatmap representation illustrating mRNA expression of the Broad Institute curated database p53 upstream regulating genes (left) and p53 downstream target genes (right) from RNA-sequencing data showing non-treated versus gemcitabine treated ATRT cultures. Average expression of the full gene set is depicted at the bottom of both heatmaps.

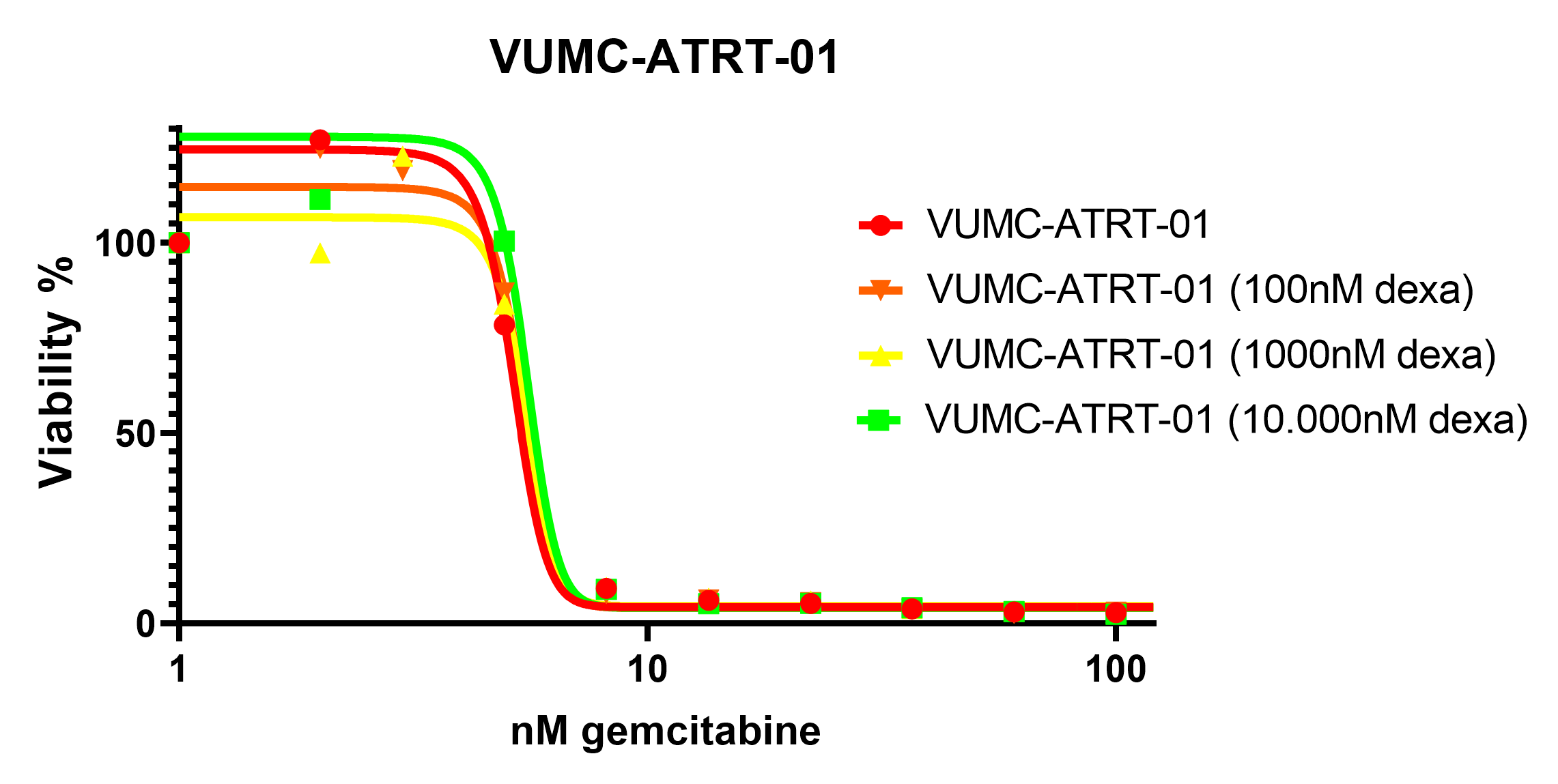

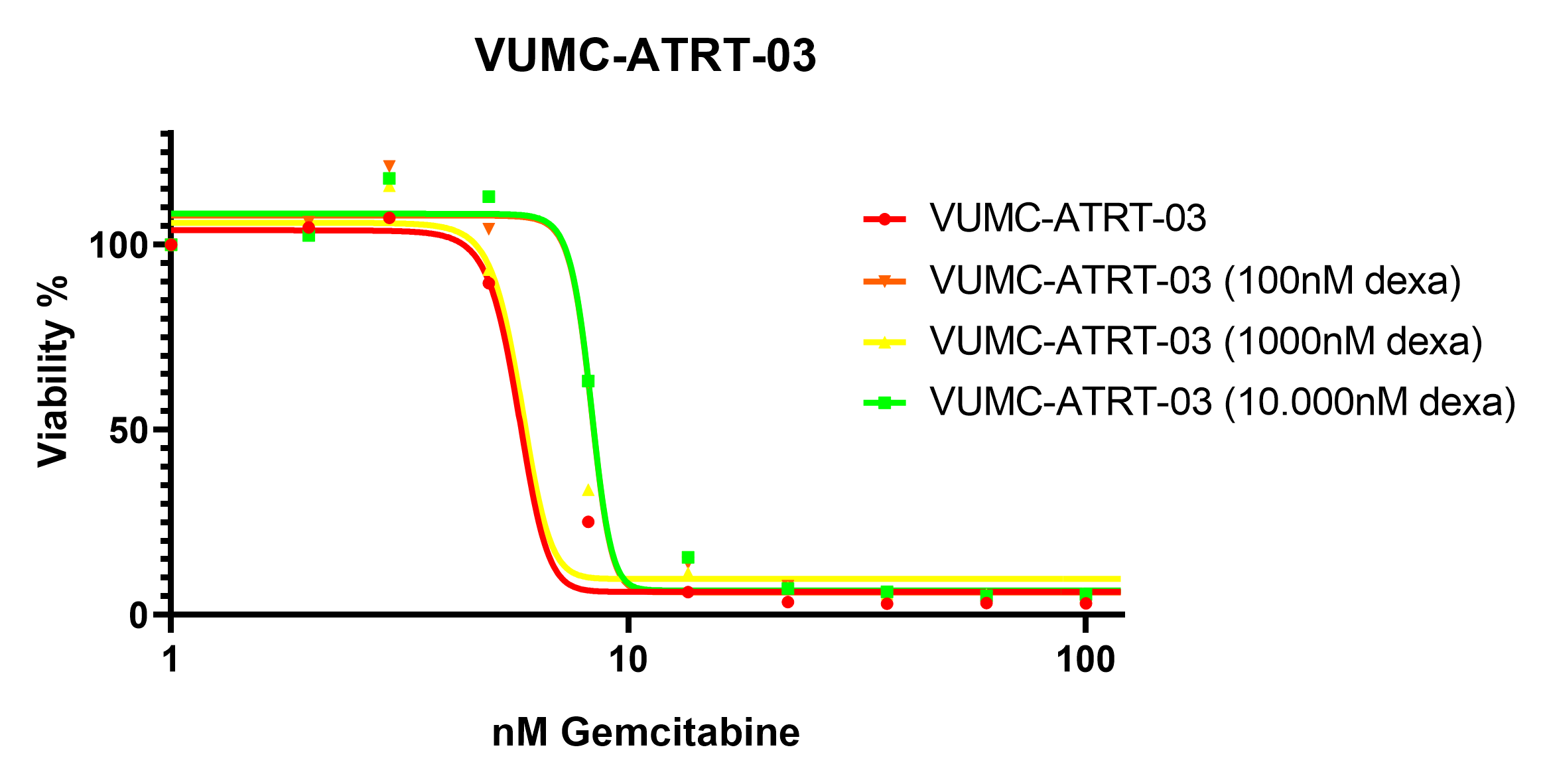

***Supplementary figure S4:*** IC50 viability curves of gemcitabine treatment combined with a dexamethasone concentration in VUMC-ATRT-01 and VUMC-ATRT-03 cells treated for 96h.

**Supplementary Tables**

| **shRNA** | **Full Hairpin Sequence** | **Sense Sequence** | **Vector** |
| --- | --- | --- | --- |
| shSIRT1 #1 | CCGGGCAAAGCCTTTCTGAATCTATCTCGAGATAGATTCAGAAAGGCTTTGCTTTTT | GCAAAGCCTTTCTGAATCTAT | pLKO.1 |
| shSIRT1 #2 | CCGGCCTCGAACAATTCTTAAAGATCTCGAGATCTTTAAGAATTGTTCGAGGTTTTT | CCTCGAACAATTCTTAAAGAT | pLKO.1 |
| shSIRT1 #3 | CCGGGCGGGAATCCAAAGGATAATTCTCGAGAATTATCCTTTGGATTCCCGCTTTTT | GCGGGAATCCAAAGGATAATT | pLKO.1 |

***Supplementary table S1:*** *shRNA sequences (sense in blue, antisense in red) of constructs used in this study.*

| **Primer** | **Sequence** |
| --- | --- |
| TP53 exon4 FWD primer | cacc**GAAGGGACAGAAGATGACAG** |
| TP53 exon4 REV primer | aaacCTGTCATCTTCTGTCCCTTC |
| hSpCas9 U6 Seq FWD | GAGGGCCTATTTCCCATGATTCC |

***Supplementary table S2:*** *Relevant sequences for cloning p53 sgRNA into vector backbone, including sequencing primer for validation. Actual TP53 guide sequence is highlighted in blue.*

| **Geneset** | **Gemcitabine > None** |
| --- | --- |
| REACTOME_HEDGEHOG_OFF_STATE (113) | -6.115 |
| REACTOME_SIGNALING_BY_HEDGEHOG (150) | -5.750 |
| REACTOME_HEDGEHOG_ON_STATE (86) | -4.574 |
| REACTOME_HEDGEHOG_LIGAND_BIOGENESIS (65) | -4.552 |
| PID_HEDGEHOG_GLI_PATHWAY (48) | -3.118 |
| *Significance* | *< -2.580* |

***Supplementary table S3:*** *Parametric assessment of geneset enrichment (PAGE) of VUMC-ATRT-03, VUMC-DIPG-10, VUMC-DIPG-11, VUMC-HGG-09, JHH-DIPG-01, HSJD-DIPG-07, and SU-pcGBM-2 cells after gemcitabine treatment (5nM, 24h), compared to non-treated conditions. All major Reactome (*[*https://reactome.org/*](https://reactome.org/)*) and Pathway Interaction Database (PID) (*[*http://pid.nci.nih.gov/*](http://pid.nci.nih.gov/)*) SHH-signaling pathways are shown significantly downregulated in gemcitabine treated conditions. Scores <-2.580 are statistically significant (FDR, p<0.05) (*[*http://r2.amc.nl/*](http://r2.amc.nl/)*).*
